# Single-cell transcriptomic profiling in inherited retinal degeneration reveals distinct metabolic pathways in rod and cone photoreceptors

**DOI:** 10.1101/2022.08.26.505393

**Authors:** Yiyi Chen, Yujie Dong, Jie Yan, Lan Wang, Shirley Yu, Kangwei Jiao, François Paquet-Durand

**Author notes:** These authors contributed equally to this work (Yujie Dong, Jie Yan).

## Abstract

The cellular mechanisms underlying hereditary photoreceptor degeneration are still poorly understood. The aim of this study was to systematically map the transcriptional changes that occur in the degenerating mouse retina at the single cell level. To this end, we employed single cell RNA-sequencing (scRNA-seq) and retinal degeneration-1 (*rd1*) mice to profile the impact of the disease mutation on the diverse retinal cell types during early post-natal development. The transcriptome data allowed to annotate 43,979 individual cells grouped into 20 distinct retinal cell types. We further characterized cluster-specific metabolic and biological changes in individual cell types. Our results highlight Ca^2+^-signaling as relevant to hereditary photoreceptor degeneration. Though metabolic reprogramming in retina, known as ‘Warburg effect’, has been documented, further metabolic changes were noticed in *rd1* mice. Such metabolic changes in *rd1* mutation was likely regulated through mitogen-activated protein kinase (MAPK) pathway. By combining single-cell transcriptomes and immunofluorescence staining, our study revealed cell type-specific changes in gene expression, as well as interplay between Ca^2+^ induced cell death and metabolic pathways.

## 1. Introduction

Retinitis pigmentosa (RP) relates to rare, genetic disorders that cause progressive vision loss [1]. Typically, RP is characterized by a two-step process where an initial, primary degeneration of rod photoreceptors is followed by a secondary loss of cone photoreceptors, eventually leading to complete blindness [2]. The disease displays a vast genetic heterogeneity, with causative mutations identified in over 85 genes [3]. One of the most extensively studied RP animal models is the retinal degeneration 1 (*rd1*) mouse. This mouse strain carries a naturally occurring nonsense mutation in the *Pde6b* gene, encoding for the β subunit of rod cGMP-phosphodiesterase-6 (PDE6). Similar mutations in the human *PDE6B* gene have been found in RP patients [4].

The mutation-induced loss-of-function of PDE6 leads to photoreceptor cGMP accumulation [5], which activates cyclic nucleotide-gated (CNG) channels. This in turn leads to Ca^2+^ influx and, further downstream, activation of Ca^2+^-dependent calpain-type proteases. In parallel, cGMP-dependent activation of protein kinase G (PKG) is associated with activation of histone deacetylase (HDAC) and poly-ADP-ribose-polymerase (PARP) [6]. cGMP-dependent, non-apoptotic cell death appears to be a common mechanism since it was identified also in a diverse set of animal models carrying mutations in a variety of RP genes [6].

Overall, the retina is characterized by high energetic and metabolic demands [7, 8], and these demands may be exacerbated by cGMP-dependent activation of CNG-channels and PKG, which in turn may be linked to the depletion of adenosine 5′-triphosphate (ATP) and nicotinamide adenine dinucleotide (NAD^+^). Notably, cGMP-dependent Ca^2+^-influx increases the demand for ATP-dependent Ca^2+^ extrusion [9], while PARP uses NAD^+^ to generate poly-ADP-ribose polymer, making it one of the main consumers of NAD^+^ [10]. Thus, ultimately RD-disease mutations may cause an energetic collapse of the photoreceptor cell [11].

Therefore, insights into the energy metabolism of the retina are crucial for understanding pathology and devising treatments for retinal diseases. Here, we took advantage of single-cell RNA sequencing (scRNA-seq) to investigate the distinct metabolic pathways underlying retinal degeneration in rod and cone photoreceptors. Our analysis suggests that a Ca^2+^-induced activation of the MAPK pathway enhances oxidative phosphorylation (OXPHOS) in *rd1* rod photoreceptors. At the same time *rd1* cones may decrease glycolytic activity. These findings may have major ramifications for future therapy developments.

## 2. Results

### 2.1. Morphologic changes in *rd1* retina

The *Pde6b* mutation in the *rd1* mouse initially affects rod photoreceptors in the outer nuclear layer (ONL). Immunofluorescence staining for PDE6B confirmed the loss of protein expression in the outer segment of *rd1* rods when compared to wild-type (WT) (Figure 1a). Concomitantly, a strong reduction of ONL thickness was observed in *rd1* retina, while the TUNEL assay revealed a large number of dying cells in the *rd1* ONL from post-natal day (P) 11 onwards (Figure 1b; quantifications in Figure 1d, e). Cone photoreceptors, while free of the disease-causing mutation, suffer from a secondary degeneration at a slower rate [12]. Immunofluorescence staining for cone arrestin was used to assess the number of surviving cone cells (Figure 1c; quantifications in Figure 1f) and showed a partial loss of *rd1* cones at P17.

**Figure 1.**
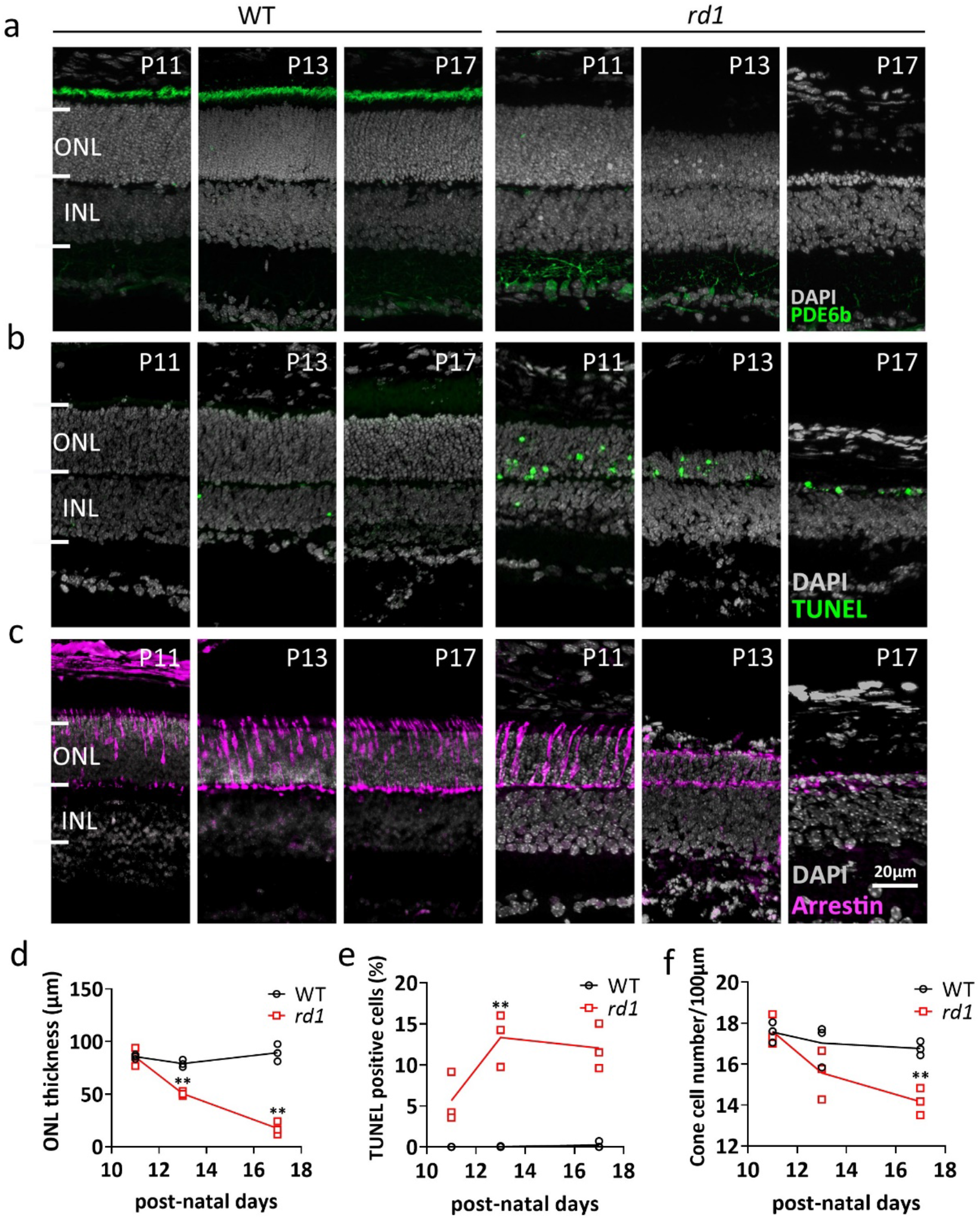
Comparison of WT and *rd1* retina during the 2^nd^ post-natal week. Retinal cross-sections were obtained at post-natal days (P) 11, 13, and 17 from wild-type (WT) and retinal degeneration 1 (*rd1*) mice. (**a**) Immunofluorescent staining (IF) for PDE6B (green), labelling photoreceptor outer segments. (**b**) TUNEL assay (green) showing dying cells in the outer nuclear layer (ONL). (**c**) IF for cone arrestin (magenta) in the ONL. (**d**) Quantification of ONL thickness, (**e**) TUNEL positive cells, and (**f**) cone cell number. Data from n=3 animals per group, expressed as mean ± SD. Statistical significance was assessed Student’s *t*-test; significance levels were: * p < 0.05, ** p < 0.01 and *** p < 0.001. DAPI (grey) was employed as nuclear counterstain. INL, inner nuclear layer.

### 2.2. Single-cell RNA sequencing analysis yields 20 clusters corresponding to eight retinal cell types

Using a droplet-based single-cell RNA-seq (scRNA-seq) platform (10×Genomics), single-cell transcriptome analysis was performed on retinal tissues from WT and *rd1* mice at post-natal day 11, 13, and 17. Sequencing data were collected from a total of 43,979 cells, of which 23,068 cells (52%) were from WT retinas and 20,911 cells (48%) from *rd1* retinas. For unbiased cell classification, we performed a t-stochastic neighbor embedding (tSNE) analysis using Seurat and identified 20 cell clusters, which were further categorized into eight distinct cell types (Figure 2a, c). These were annotated as follows: rod photoreceptors (clusters 0, 1, 2, 3), cone photoreceptors (cluster 9), bipolar cells (clusters 5, 6, 7, 10-14), amacrine cells (clusters 8, 16), Müller cells (clusters 4, 19), horizonal cells (cluster 15), microglia cells (cluster 17), and vascular cells (cluster 18). Clusters corresponding to retinal pigment epithelium (RPE) and retinal ganglion cells were not detected in this study, probably due to their relatively low number in the total retinal cell populations. The abundance of WT *vs. rd1* cells at each time point is illustrated in Figure 2b, while the relative abundance of cell types in the WT and *rd1* cell population is represented in Figure 2d.

**Figure 2.**
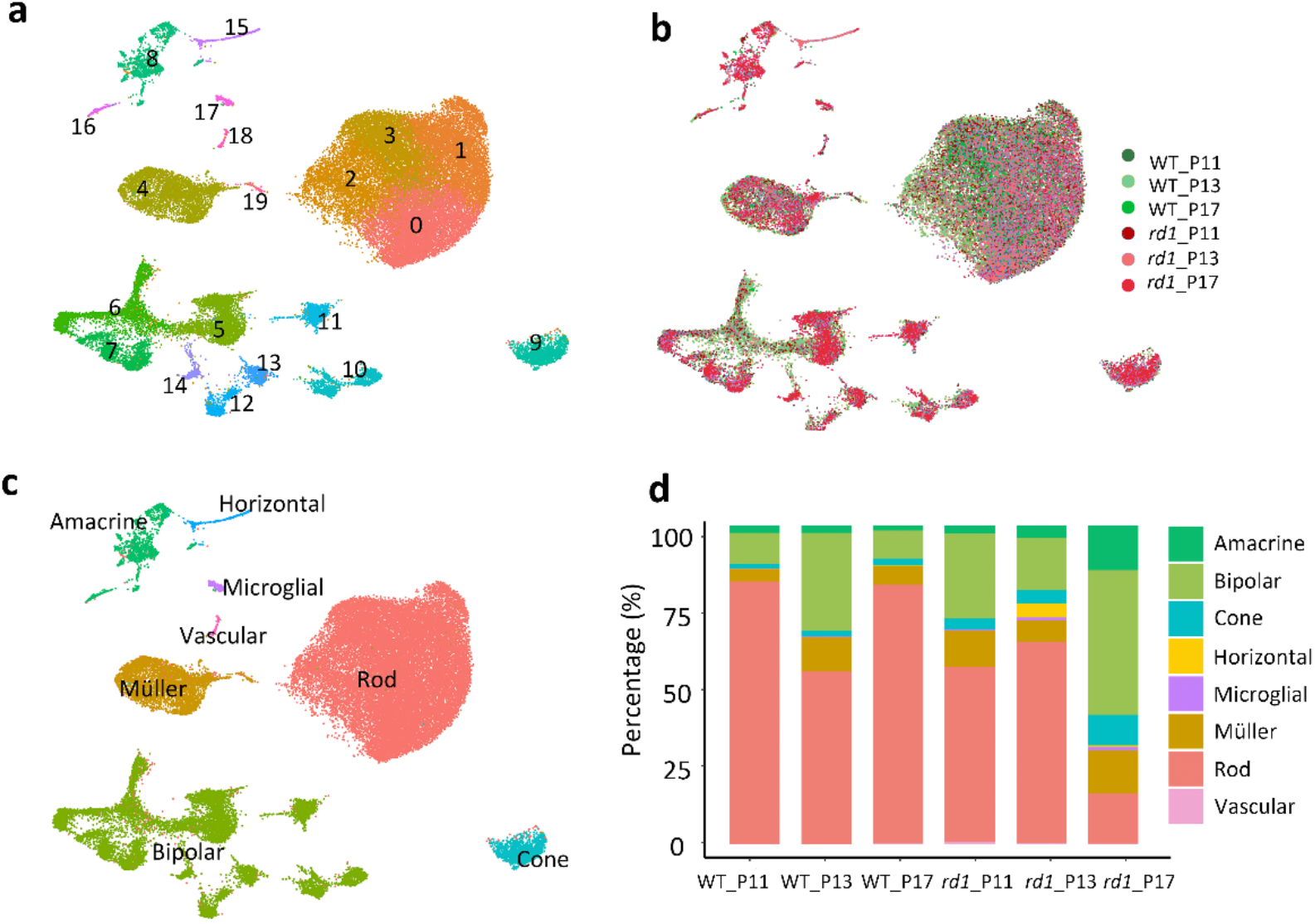
Single-cell RNAseq analysis of WT *vs. rd1* mouse retinas. (**a**) A total of 43,979 cells isolated from wild-type (WT) and retinal degeneration-1 (*rd1*) retinas were grouped into 20 identified clusters. (**b**) Distribution of all isolated cells clustered temporally. (**c**) All isolated cells color-coded by retinal cell type. (**d**) Bar graphs displaying the percentages of cell types identified in WT and *rd1* samples at P11, P13, and P17. Note the marked decrease of rod photoreceptors in the *rd1* P17 samples.

We then identified the top 35 most regulated genes across the eight most prominent cell types, each of which was characterized by a set of differentially expressed genes (Figure 3). For instance, the genes *Rho, Nr2e3, Nrl, Pdc*, and *Rp1* showed the highest expression on rod, while *Opsn1sw, Opn1mw, Pde6h, Arr3*, and *Gnat2* were expressed in cones. Genes up-regulated in Müller cells included *Zfp36l1, Dbi, Apoe, Slc1a3, Sparc*, and *Pcp2*. The *Pcp2, Trpm1, Isl1, Grm6*, and *Trnp1* genes were prominent in bipolar cells. The genes *Meg3, C1ql1, Snhg11, Tfap2b*, and *Pcsk1n* were enriched in amacrine cells. *C1ql1* and *Snhg11* were also highly expressed in horizontal cells, as well as *Tfap2b, Psk1n, Scl4a3, Calb1, Sept4*, and *Tpm3*. The genes *Ctsd, Ccl4, C1qb*, and *C1qc* ranked high in microglial cells. *Pcp2, Trpm1, Meg3*, and *Lgfbp7* were highly expressed in vascular cells. Additional data for the 20 identified clusters, including the top four markers for each cluster, are presented in Supplementary Figure S1.

**Figure 3.**
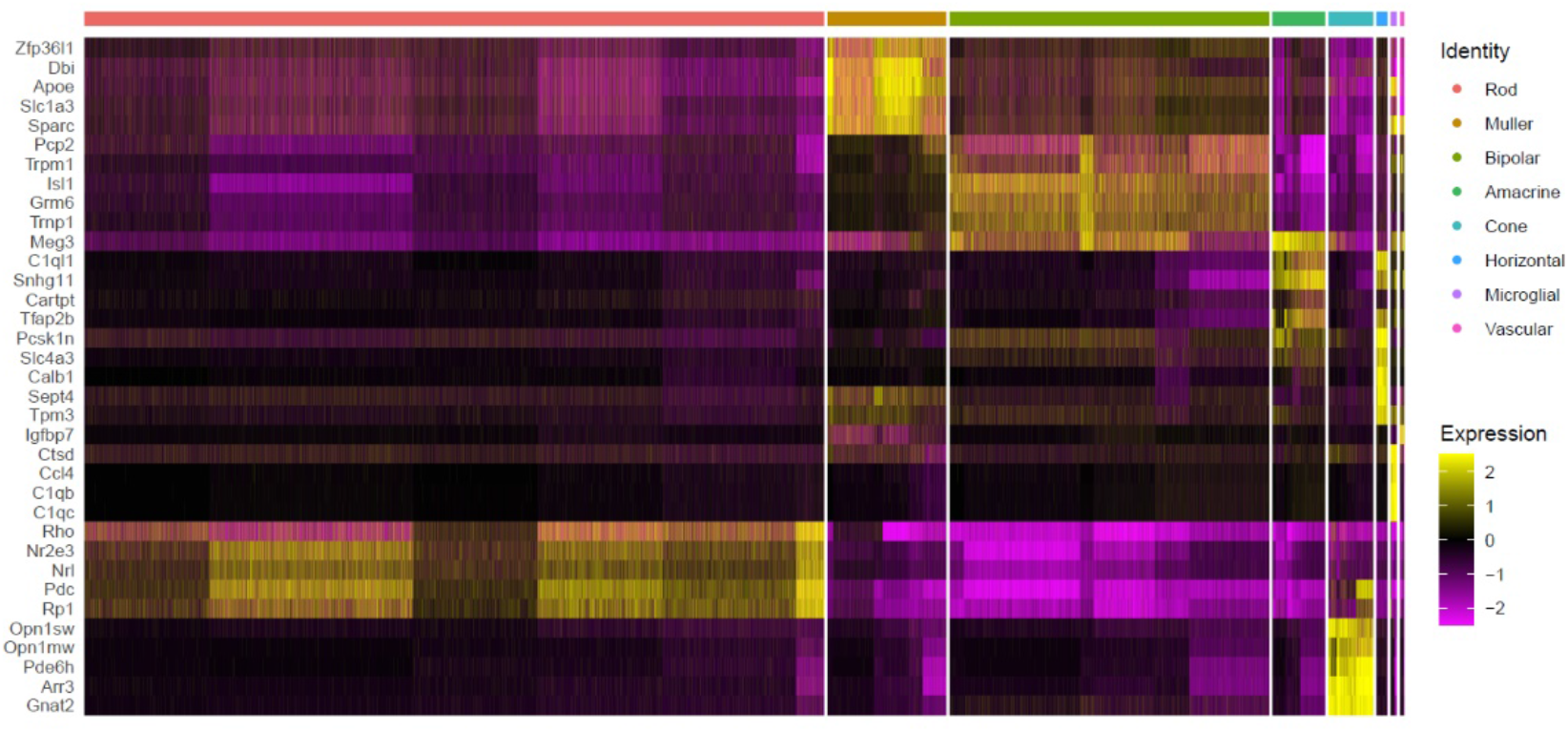
Expression of 35 marker markers across eight cell types in WT and *rd1* mouse. The Seurat FindMaker function was used to analyze differences between cell populations and to screen each population for genes with an average log2 fold-change (FC) > 1.5 and a *p*-value < 0.05 relative to all other cells. The heat map displays expression of top 35 enriched genes across the eight most abundant cell types. Cell types are arranged on x-axis and genes are organized along y-axis. A more yellow color indicates that a cell population highly and specifically expresses a given gene, magenta indicates low gene expression.

### 2.3. Differential gene expression analysis in WT and *rd1* retinal cell types

An analysis of differentially expressed genes (DEGs) was first carried out on rods and cones at corresponding ages (Supplemental dataset 1). A volcano plot of DEGs in *rd1* retina at P13 showed 151 up- and 111 down-regulated genes compared to WT (Figure 4a). Functional pathways and networks were analyzed for these DEGs with log2 fold-change (FC) greater than 0.5. This was followed by a gene ontology (GO) enrichment analysis, which indicated that biological processes (BP) related to visual perception, phototransduction, retinal development, and apoptosis were prominently regulated in *rd1* rod photoreceptors (Figure 4a). The ‘cellular response to Ca^2+^ ions’ was also among the Top 10 most regulated pathways, indicating that Ca^2+^ played an important role in the degeneration of photoreceptors. Functional enrichment and interactome analysis using the Metascape web portal [13] produced an interactome network in which each enriched term was represented as a node and where connections between nodes were plotted for Kappa similarities over 0.3 across all 262 gene candidates. An enrichment of molecular pathways similar to the GO analysis was also found in the KEGG database, where phototransduction, neurotransmitter transport, and synaptic function were among the most regulated processes.

**Figure 4.**
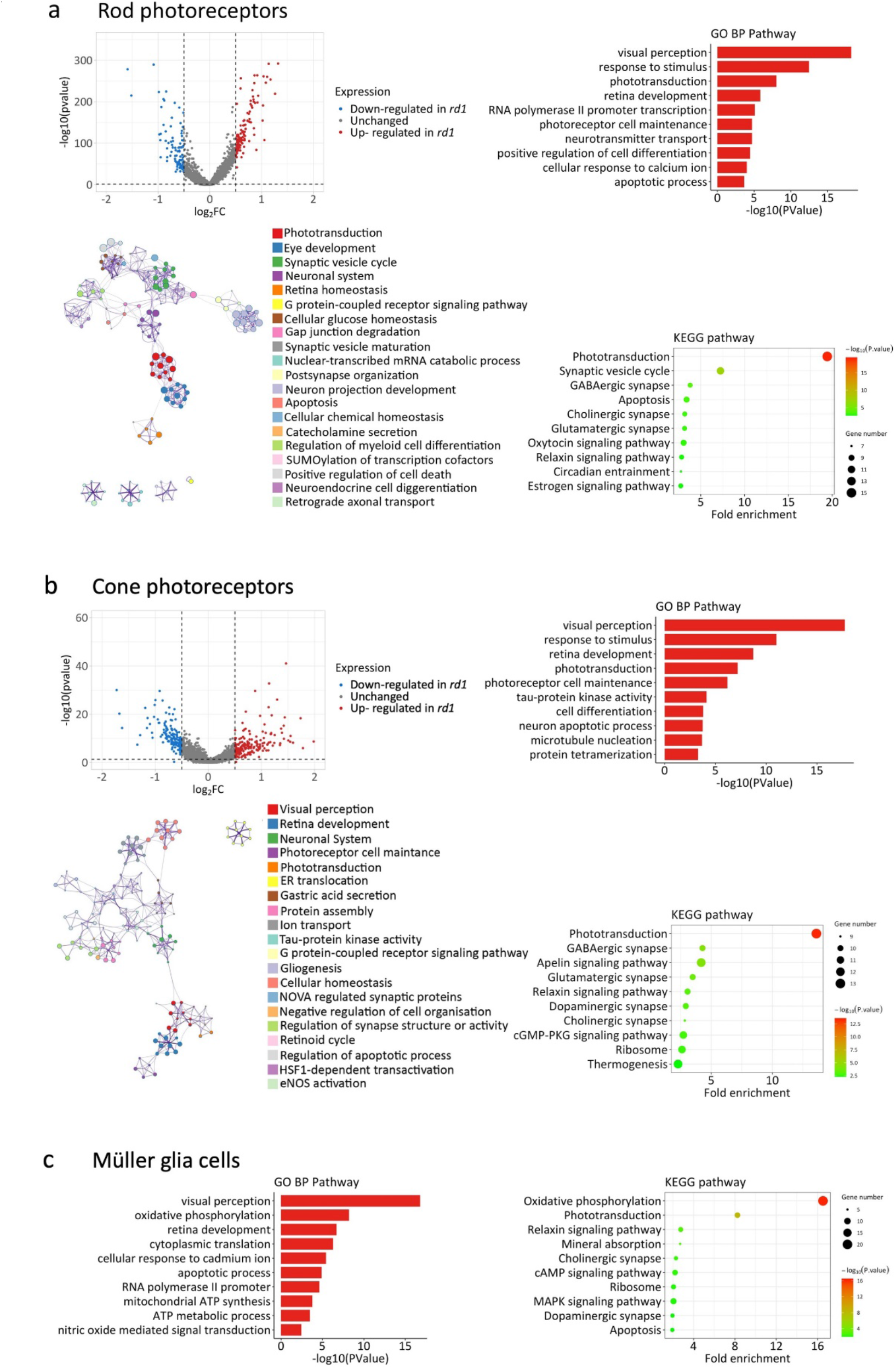
Network and pathway analysis for rods, cones, and Müller glia cells. Data showing differentially enriched genes (DEGs) in *rd1* retinal cell types, compared to wild-type (WT). (**a**) Network and pathways changes in rod photoreceptors at post-natal day (P)13, *i*.*e*. at the peak of rod degeneration. Top left panel: Volcano plot of all genes significantly regulated in *rd1* rods (log2 fold-change (FC) threshold = 0.5). Blue dots for down-regulated genes, red dots for up-regulated genes, grey indicates no significant change between *rd1* and WT. Top right: Gene ontology (GO) biological category (BP) analysis of DEGs showing top 10 most enriched GO BP terms. Bottom left: Metascape biological network analysis with each term represented by a circle node, size proportional to the number of input genes falling under that term, and color representing cluster identity. Terms with a similarity score > 0.3 are linked by an edge. Term labels were only shown for one term per cluster for clarity. Bottom right: KEGG pathway analysis of DEGs showing top 10 enriched terms. (**b**) Network and pathways changes in cone photoreceptors at P17, *i*.*e*. at the start of cone degeneration. Top left: Volcano plot of all genes significantly regulated in *rd1* cones. Blue dots for down-regulated genes, red dots for up-regulated genes, grey indicates no significant change between *rd1* and WT. Top right: Gene ontology (GO) biological category (BP) analysis of DEGs showing top 10 most enriched GO BP terms. Bottom left: Metascape biological network analysis. Bottom right: KEGG pathway analysis showing top 10 enriched terms. (**c**) Pathways analysis on Müller cells at P13. Left: Top 10 enriched pathways in gene ontology biological process analysis. Right: KEGG pathway analysis showing top 10 enriched terms.

Since cone photoreceptors were affected by the degeneration later than rods (*cf*. Figure 1c) the corresponding cone pathway and network analysis was performed for the P17 time-point (Figure 4b). A volcano plot for DEGs in P17 cones revealed 219 up-genes and 154 down-regulated genes.

In both GO and KEGG based analyses, the pathways most altered in cones were related to visual perception, retinal development, phototransduction, and synaptic functions. Additionally, tau-protein kinase activity, microtubule nucleation, and protein tetramerization were unique to cone cells as identified by GO pathway analysis. Interestingly, the cGMP-PKG signaling pathway was also related to cone degeneration at P17. Metascape network analysis revealed, among other things, a close interaction between visual perception, retinal development, phototransduction, and photoreceptor cell maintenance.

Because of their tight connection to photoreceptors, the pathway analysis was extended to Müller glial cells (Figure 4c). The GO analysis of Müller cells also revealed pathways related to visual perception and retinal development. Furthermore, oxidative phosphorylation, ATP synthesis, and nitric oxide signaling pathways were enriched. A further analysis of pathways changed in *rd1* amacrine and horizontal cells is provided in Supplemental Figure S2.

Overall, the analysis of functional pathways and networks indicated that Ca^2+^-signaling and alterations in cellular metabolism might be connected with retinal degeneration and cell death.

### 2.4. Transcriptional changes related to cell death in *rd1* mutant retina

Previous research had related *rd1* mouse photoreceptor degeneration to non-apoptotic mechanisms that involved the activity of CNG channels, PKG, HDAC, and PARP [6]. We therefore assessed the expression of corresponding genes in rods and cones, starting with that of the various PDE6 genes (Figure 5). While *Pde6b* was clearly down-regulated in *rd1* rods, interestingly, both *Pde6a* and *Pde6g* were transiently elevated at P13. *Pde6h*, which codes for the cone-specific inhibitory subunit of PDE6, was found to be highly up-regulated in cones at P13. Especially the up-regulation of the genes encoding for the inhibitory PDE6G and PDE6H subunits may indicate an attempt to compensate for the loss of PDE6B expression.

**Figure 5.**
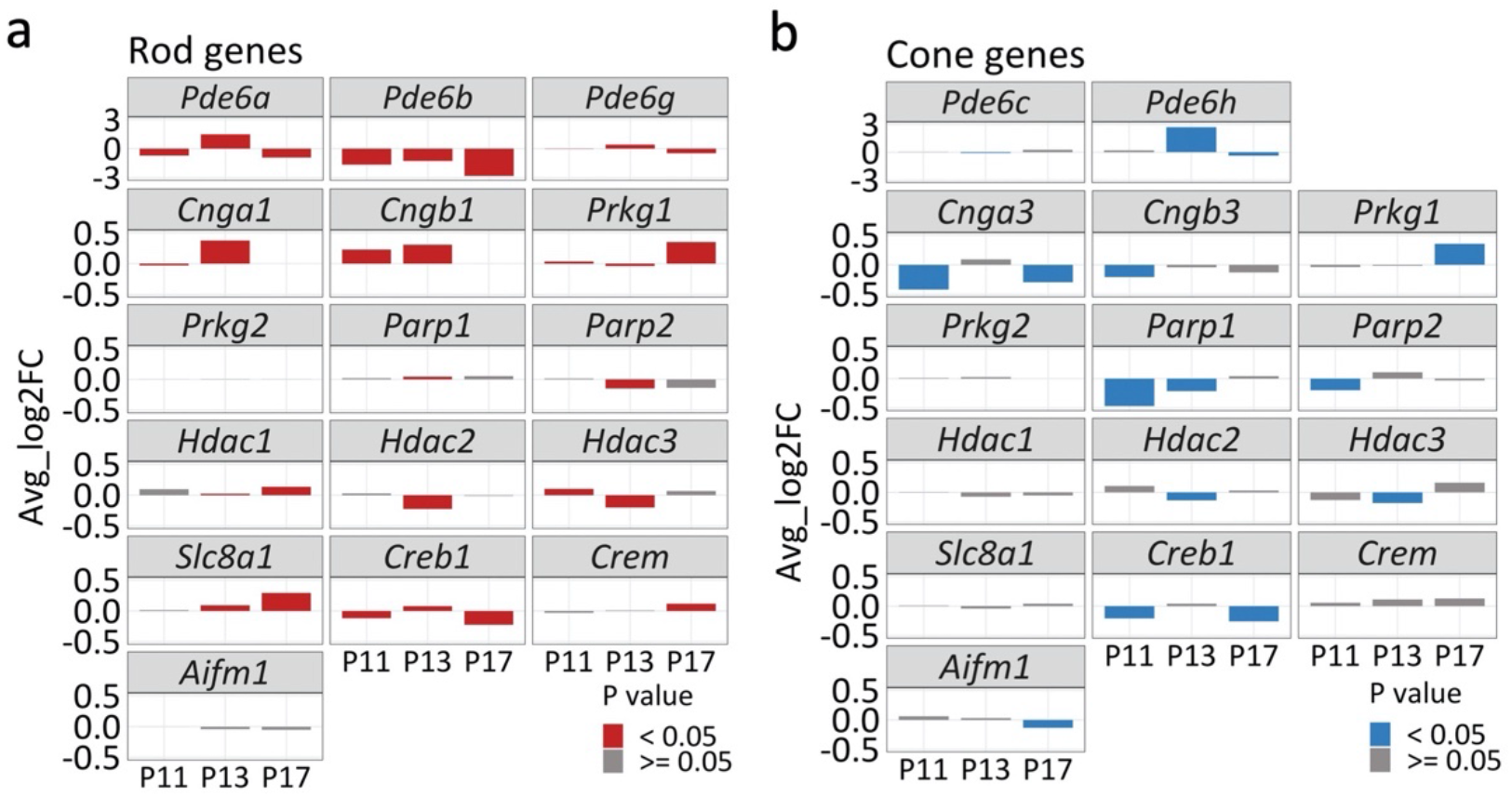
Expression of genes related to cell death in rods and cones. Bar plots showing transcriptomic changes in genes coding for phosphodiesterase-6 (PDE6; *i*.*e*., *Pde6a, Pde6b, Pde6c, Pde6g, Pde6h*), cyclic-nucleotide-gated channels (CNG-channels; *i*.*e*., *Cnga1, Cngb1, Cnga3, Cngb3*), protein kinase G (PKG; *Prkg1, Prkg2*), poly(ADP-ribose) polymerase (PARP; *Parp1, Parp2*), histone deacetylase (HDAC; *Hdac1, Hdac2, Hdac3*), Na^+^/Ca^2+^ exchanger-1 (NCX1; *Slc8a1*), cAMP response element-binding protein-1 (*Creb1*) and cAMP response element modulator (*Crem*), and apoptosis inducing factor-1 (*Aifm1*). The x-axis indicates postnatal day (P), y-axis depicts average log2 fold change; positive values indicating higher expression in *rd1* rods/cones. Bars were color-coded (red for rods; blue for cones) for *p*-values < 0.05.

Whatever the case, loss of PDE6 activity produces an accumulation of cGMP within photoreceptors [14, 15], keeping CNG channels open. The rod CNG channel is composed of three CNGA1 subunits and one CNGB1 subunit [16]. From our results, both *Cnga1* and *Cngb1* were significantly upregulated at P11 and P13, but strongly reduced at P17. *Cnga3* and *Cngb3*, which encode the α- and β-subunit of the CNG channel in cones, were both down-regulated at P11 and P17.

PKG is another key effector of cGMP-signaling and encoded by the *Prkg1* and *Prkg2* genes [17]. *Prkg1* transcription was upregulated at P17 in both rods and cones, while *Prkg2* was not significantly changed.

Downstream of CNG channel and PKG activity, PARP and HDAC have been shown to contribute to *rd1* photoreceptor cell death [18]. We found both *Parp1* and *Parp2* to be downregulated, especially on cones. In rods, *Hdac1* was found to be significantly increased at P17, indicating a possible role in the final phase of rod cell death. Both *Hdac2* and *Hdac3* were essentially downregulated in *rd1* rods and cones.

Apart from CNG channels, an important regulator of intracellular Ca^2+^-levels is the Na^+^/Ca^2+^ exchanger-1 (NCX1) encoded for by the *Slc8a1* gene. This gene was increasingly upregulated in rods at P13 and P17 but remained unchanged on cones.

The CREB1 and CREM transcription factors together function as a central hub that regulates transcription in response to various stressors, metabolic changes, and developmental signals [19, 20]. Importantly, they are targets for PKG phosphorylation and may serve as transducers of cGMP-signaling [21]. A downregulation of *Creb1* was seen both in rods and cones at P11 and P17, whereas *Crem* levels were only increased in rods at P17. We further performed IF on CREB1 and CREM on WT and rd1 retina, at P11, P13, and P17, albeit without finding obvious changes in protein expression (Supplemental Figure S3).

Surprisingly, the gene *Aifm1* which encodes the mitochondrial protein apoptosis inducing factor (AIF), an important regulator of programmed cell death [22], did not exhibit any significant transcriptional changes in rods. However, a down-regulation of the *Aifm1* gene was observed in cone photoreceptors at P17. IF for AIF protein strongly labelled mitochondria-containing structures, such as the photoreceptor inner segments and synapses. However, no significant changes in AIF protein expression were seen between WT and *rd1* samples (Supplemental Figure S3).

### 2.5 Photoreceptors in *rd1* mutant retina undergo a metabolic switch

Several previous studies indicated a disturbance in energy metabolism during photoreceptor degeneration [23, 24], prompting us to examine transcriptional changes on related pathways.

Using gene set enrichment analysis (GSEA) [25] and the molecular signatures data base (MSigDB) gene sets, we investigated ‘glycolysis’, ‘tricarboxylic acid (TCA) cycle’ and ‘oxidative phosphorylation (OXPHOS)’, as well the cell death related pathways ‘apoptosis’ and ‘DNA repair’. Between P11 and P17, all five pathways displayed significant changes in rods and/or cones (Figure 6a). Notably in rods, the energy metabolism related pathways TCA cycle, OXPHOS, and glycolysis were significantly altered.

**Figure 6.**
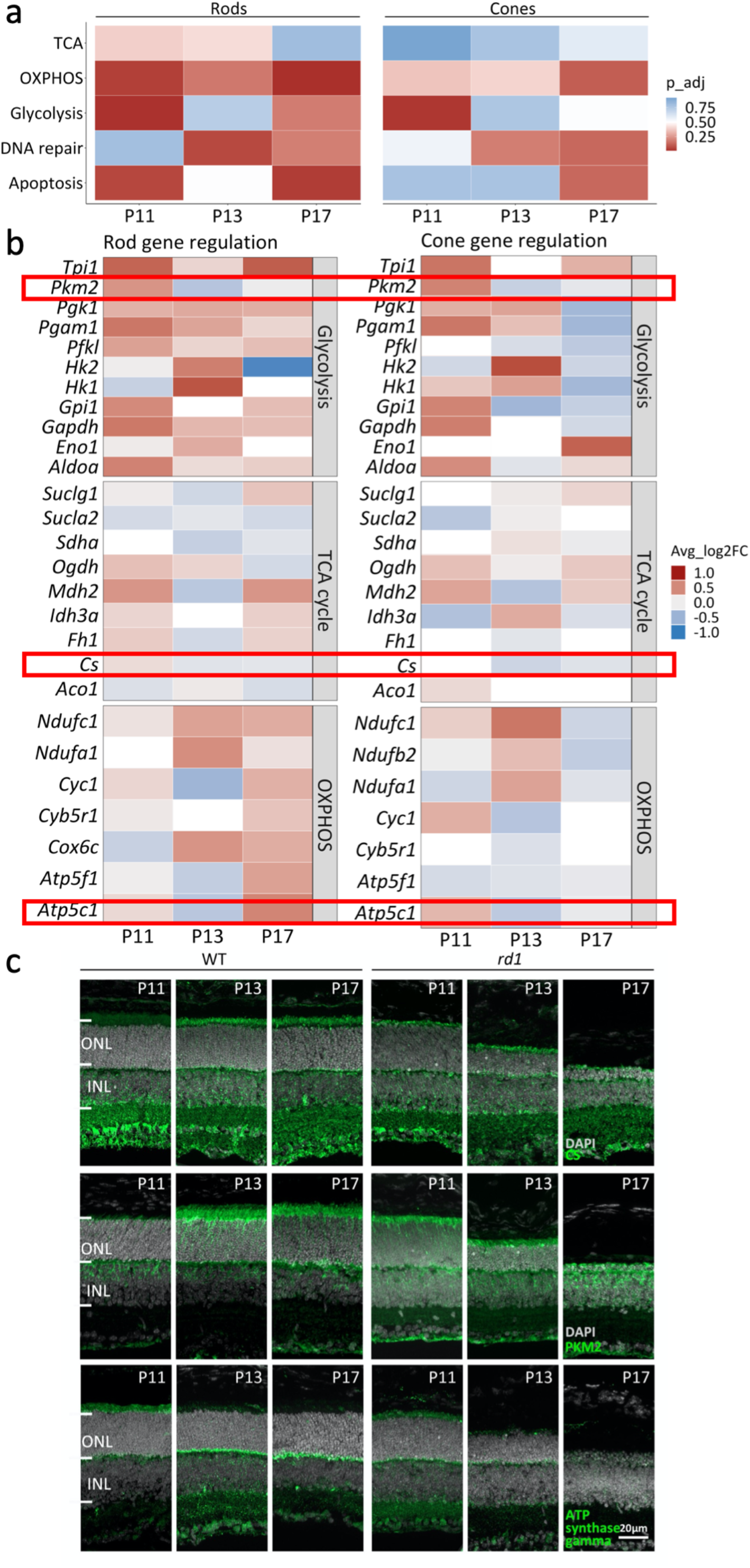
Analysis of genes and enzymes of the TCA cycle, OXPHOS, and glycolysis. (**a**) Gene set enrichment analysis (GSEA) of differentially expressed genes (DEGs) performed on retinal degeneration-1 (*rd1*) *vs*. wild-type (WT) retina. Heatmap showing rod and cone DEGs differentially enriched from post-natal (P) day 11 to P17, in five pathways: Tricarboxylic acid (TCA) cycle, oxidative phosphorylation (OXPHOS), glycolysis, DNA repair, and apoptosis. (**b**) Heatmap showing gene expression changes in individual genes involved in glycolysis, TCA cycle, and OXPHOS. Key enzymes highlighted by red frames. Gene squares filled in white when p value > 0.5. (**c**) Immunofluorescence (IF) staining for citrate synthase (CS), pyruvate kinase (PKM2), and ATP synthase gamma. DAPI (grey) was employed as nuclear counterstain. ONL, outer nuclear layer; INL, inner nuclear layer.

To determine how energy metabolism may have changed during *rd1* degeneration, we examined sets of individual genes corresponding to each of these three pathways in more detail (Figure 6b). Remarkably, most genes related to glycolysis were strongly upregulated in rods between P11 and P17, while in cones glycolysis genes were down-regulated at P17. No clear trend for regulation of TCA cycle related genes was apparent for either rods or cones. However, genes related to OXPHOS were strongly upregulated in rods at P17, while they were strongly down-regulated in cones at the same time-point. These results indicated a switch in metabolism towards an increased ATP-production in rods, perhaps to adapt to increasing demand caused by degenerative processes. In cones, however, the down regulation of energy metabolism related genes suggested a decreased ATP-production at P17, *i*.*e*. at the onset of cone degeneration.

To verify the observed transcriptomic regulation, we performed immunofluorescence staining (IF) for key enzymes of the TCA cycle, OXPHOS, and glycolysis. Citrate synthase (CS) catalyzes the first step of the TCA cycle. As shown by IF, CS was expressed strongly in photoreceptor inner segments and synapses, and in *rd1* retina CS protein expression appeared to weaken as the degeneration progressed from P11 to P17 (Figure 6c). Pyruvate kinase M2 (PKM2) is a glycolytic enzyme that catalyzes the conversion of phosphoenolpyruvate to pyruvate. PKM2 IF was found on photoreceptor inner segments and its expression also seemed to decrease during *rd1* degeneration. ATP synthase gamma is a subunit of ATP synthase, a critical enzyme for oxidative phosphorylation and mitochondrial ATP production [26]. It is encoded for by the *Atp5c1* gene. ATP synthase gamma was localized prominently on photoreceptor synapses, and in *rd1* retina its expression decreased strongly from P11 to P17.

### 2.6 The MAPK signaling pathway coordinates energy metabolism and cell death

The mitogen-activated protein kinase (MAPK) pathway is Ca^2+^-dependent [27] and has been implicated in controlling cellular metabolism [28]. Hence, the MAPK pathway could potentially serve as an intermediary between excessive cGMP-signaling and alterations in energy metabolism. We therefore assessed transcriptional changes in MAPK-pathway related genes, identifying significant changes in 46 rod and six cone genes (Figure 7).

**Figure 7.**
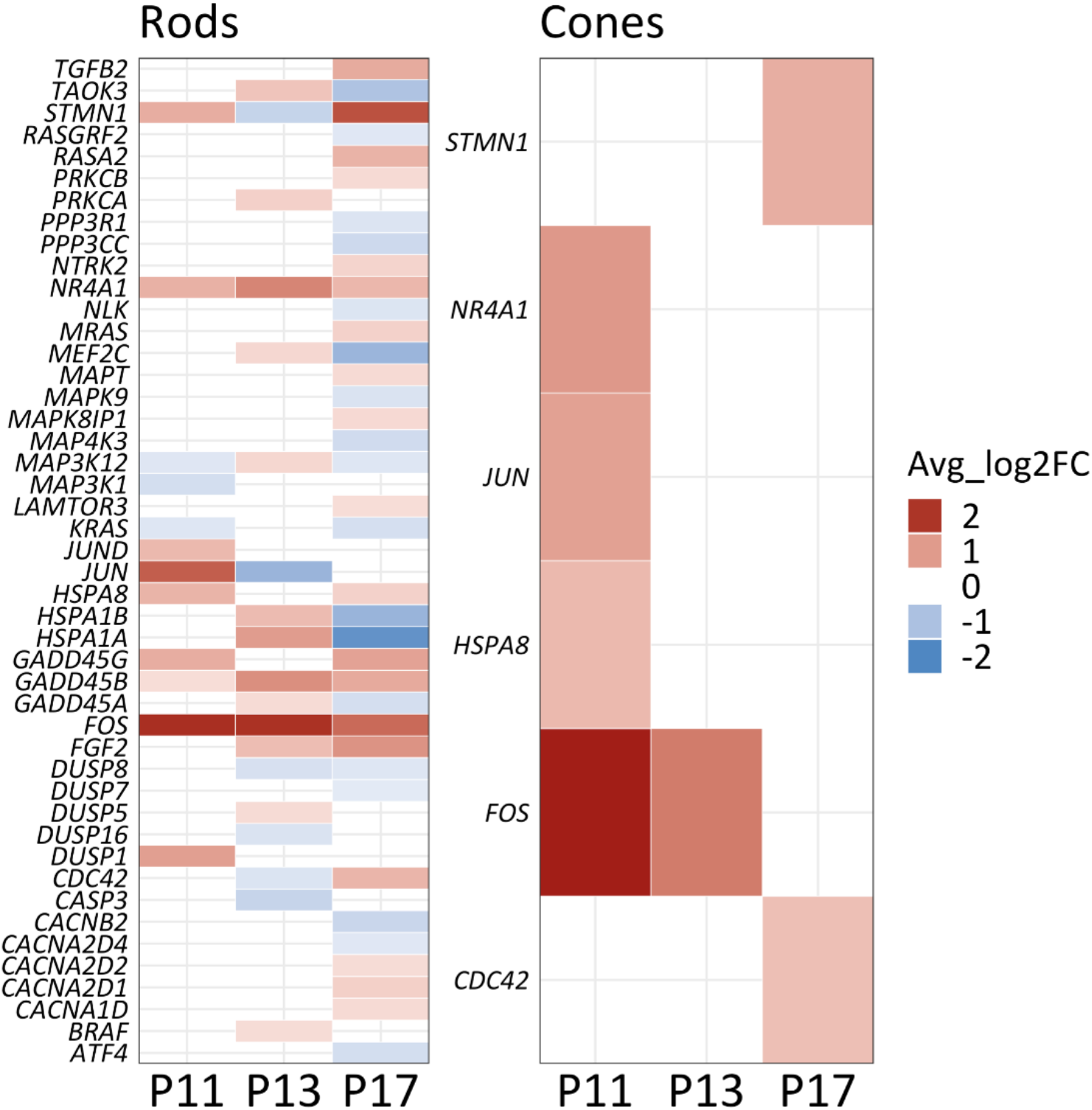
Transcriptional changes in MAPK-pathway related genes. Heatmap showing genes related to the MAPK gene set (MSigDB, gene set enrichment analysis) differentially expressed between wild-type and *rd1* rod and cone photoreceptors (*p*-value ≤ 0.5; average log2 fold-change > 0.3) for the post-natal (P) day 11, P13, and P17 time-points. Blue indicates down-regulated genes, red indicates up-regulation. A total of 46 MAPK pathway genes were significantly altered in rods, whereas six genes were altered in cones.

Activation of the MAPK-signaling pathway typically follows a three-tier kinase module in which a MAP3K phosphorylates and activates a MAP2K, which in turn phosphorylates and activates a MAPK [29]. Among the significantly changed genes, *Braf, Taok3, Nlk, Map3k1, Map3k12, Map4k3*, and *Mapk9* belong to the three layers of activating kinases. Additionally, *Fos, Nr4a1, Jun, Jund*, and *Atf4* are transcription factors involved in MAPK signaling. A significant regulation was also observed for a group of genes that code for voltage-gated Ca^2+^-channel (VGCC) subunits, including *Cacna1d, Cacnb2, Cacna2d1, Cacna2d2*, and *Cacna2d4*. This could imply that photoreceptor Ca^2+^-levels may be influenced not only by CNG-channel activity but also by VGCCs as suggested also by recent pharmacological studies [30, 31]. Note that of the six genes significantly regulated in cone photoreceptors five also appear as regulated in rods (Figure 7). At any rate, our data suggests the MAPK pathway as an important regulator of intracellular signaling during rod and cone degeneration and consequently also as a potential target for therapeutic interventions in RD-type diseases.

## 3. Discussion

In this study, we have combined scRNA-seq, immunofluorescence, and cell death detection to gauge the molecular pathways underlying *rd1* phenotype. The scRNA-seq dataset annotated 43,979 individual cells grouped into eight distinct retinal cell types and confirmed the key role of cGMP- and Ca^2+^-signaling pathways in *rd1* photoreceptor degeneration. Molecular profiling of rods and cones indicated a shift from glycolysis towards TCA-cycle and OXPHOS activity, likely reflecting increased energy demand. Moreover, our analysis suggested that the MAPK pathway may act as an intermediary between cGMP- and Ca^2+^-signaling on the one hand and cellular metabolism on the other hand.

### 3.1. cGMP- and Ca^2+^-signaling in *rd1* retinal degeneration

The loss of rod photoreceptors in *rd1* mice was previously found to depend on a non-apoptotic cell death mechanism triggered by high intracellular cGMP-levels [6, 32]. In photoreceptors high cGMP likely activates the prototypic effectors PKG and CNG channels, leading, among other things, to increased Na^+^- and Ca^2+^-influx [33, 34]. The continuous depolarization effected by CNG-channel activity causes a sustained activation of VGCCs leading to more Ca^2+^ influx [30, 35]. The Na^+^/Ca^2+^ exchanger (NCX) type antiporters encoded by the *Slc8* gene family utilize the Na^+^ gradient to extrude Ca^2+^ [36, 37]. The transcriptional upregulation of *Slc8a1* may represent an attempt to counterbalance Ca^2+^ overload. However, the Na^+^ gradient required for NCX to function is established to a large extent by the ATP-driven Na+/K+ exchanger (NKX), so that NCX-dependent Ca^2+^-extrusion will likely place an additional burden on photoreceptor energy metabolism[38]. The link between excessive cGMP-signaling, PKG activity, and cell death has been well established for the *rd1* mouse and other RD animal models [6, 14, 39]. However, at present it is still unclear which PKG isoform may be responsible for photoreceptor death. Here, we observed an up-regulation of *Prkg1* but not *Prkg2* in rod photoreceptors during the critical degeneration phase. Remarkably, in cones increased *Prkg1* gene transcription was observed only at P17, *i*.*e*. at the beginning of cone degeneration [40], indicating that also cone death may be triggered by *Prkg1* over-activation.

In retinal degeneration PKG activity is associated with an over-activation of PARP and HDAC [30, 41, 42]. PARPs are a superfamily of ADP-ribosylating enzymes and are known to play important roles in DNA repair and the maintenance of genome integrity [43-45]. The main function of HDACs consists in removing acetyl groups from DNA-binding histone proteins, which is generally associated with a decrease in chromatin accessibility for transcription factors and hence represses gene expression [46]. Though *Parp1* and *Parp2* as well as *Hdac2* and *Hdac3* did not show significant changes in P17 rods, it is important to note that activity of PARP and HDAC relays mostly on post-transcriptional regulation and may therefore not be adequately resolved with transcription-based analysis [6, 47].

### 3.2 Metabolic responses of degeneration retina

The retina and especially the photoreceptors are characterized by a very high energy expenditure, most of which may be due to the active transport of ions against their concentration and electrical gradients [8]. In the *rd1* condition, higher levels of Ca^2+^-influx entail increased ATP-consumption for Ca^2+^ extrusion [11, 32]. Besides, increased PARP activity consumes large amounts of NAD^+^ [48] and low levels of this critical electron acceptor may further impair mitochondrial ATP production [49].

Paradoxically, even in the presence of oxygen the retina converts a large fraction of its available glucose to lactate rather than oxidizing it completely to carbon dioxide. This phenomenon was first observed by Otto Warburg and is known as ‘aerobic glycolysis’ or ‘Warburg effect’ [50]. Per molecule of glucose aerobic glycolysis produces only two ATP molecules while the TCA cycle coupled to OXPHOS can generate up to 36 ATP molecules. Thus, from an energetic point of view mitochondrial oxidative metabolism is by for more efficient than cytosolic glycolytic metabolism.

To date it is not clear why the retina uses the seemingly inefficient aerobic glycolysis to generate ATP, although it has been speculated that glycolytic intermediates, including precursors of nucleic acids, lipids, and amino acids, may be required for retinal anabolic activity [51]. Perhaps even more surprising is that our study suggests that during retinal degeneration glycolytic activity may decrease while OXPHOS appears to increase. Future studies may reveal whether this constitutes an attempt to ramp up ATP-production or whether it reflects a decrease in the need for glycolytic intermediates. Moreover, our data indicates that this switch in energy metabolism may be regulated by the MAPK pathway.

### 3.3 MAPK pathway regulates crosstalk between Ca^2+^ and energy metabolism

The mitogen-activated protein kinase (MAPK) cascade has been shown to play a key role in the regulation of cell proliferation, differentiation, and death [52]. In mammals, MAPKs are divided into three subfamilies: extracellular signal-regulated kinases (ERKs), Jun-N-terminal kinases (JNKs), and p38 kinases [53]. MAPK signaling is activated by a three-tier kinase module that consists of a MAP3K phosphorylating and activating a MAP2K, which in turn activates a MAPK [29]. According to our results, ERK and JNK pathways were upregulated during retinal degeneration. While previously Ca^2+^ overload in photoreceptor leads was found to cause activation of Ca^2+^-dependent calpain-type proteases [54, 55], high Ca^2+^ may additionally activate MAPKs and ERKs directly or indirectly through protein kinase C. The MAPK signaling pathway has recently been found as a key regulator of the Warburg effect in cancer and in metabolic reprogramming [28]. Notably, the MAPK cascade promotes aerobic glycolysis through PKM2 phosphorylation, as well as regulating the expression and activity of transcription factors that directly control the expression of glycolytic enzymes, including Myc associated factor X (MAX) and c-MYC [56].

## 4. Materials and Methods

### Animals

C3H HeA *Pde6b*^*rd1*^/^*rd1*^ (*rd1*) and congenic C3H HeA *Pde6b*^+/+^ wild-type (WT) mice were used, two mouse lines that were originally created in the lab of Somes Sanyal at the University of Rotterdam [57]. All efforts were made to minimize the number of animals used and their suffering, notably through the use of *in vitro* experimentation (see below) and the use of both retinae from the same animal. Animals were housed in the specified pathogen-free facility of the Tübingen Institute for Ophthalmic Research, under standard white cyclic lighting in type-2 long cages with a maximum of five adults per cage. They had free access to food and water and were used irrespective of gender. Protocols compliant with the German law on animal protection were reviewed and approved by the ‘Einrichtung fur Tierschutz, Tierärztlichen Dienst und Labortierkunde’ of the University of Tübingen (AK 02/19 M, notice acc. to §4 German law on animal protection) and were following the association for research in vision and ophthalmology (ARVO) statement for the use of animals in vision research.

### Immunofluorescence

Fixed slides were dried at 37°C for 30 min and rehydrated for 10 min in PBS at room temperature (RT; 21°C). For immunofluorescent labelling, the slides were incubated with blocking solution (10% normal goat serum, 1% bovine serum albumin in 0.3% PBS-Triton X 100) for 1 h at RT. The primary antibodies were diluted (see Table 1) in blocking solution and incubated at 4°C overnight. The slides were then washed with PBS, three times for 10 min each. Subsequently, a corresponding secondary antibody, diluted in PBS (see Table 1), was applied and incubated for 1 h at RT. Lastly, the slides were washed with PBS and covered in Vectashield with DAPI (Vector, Burlingame, CA, USA).

**Table 1.**
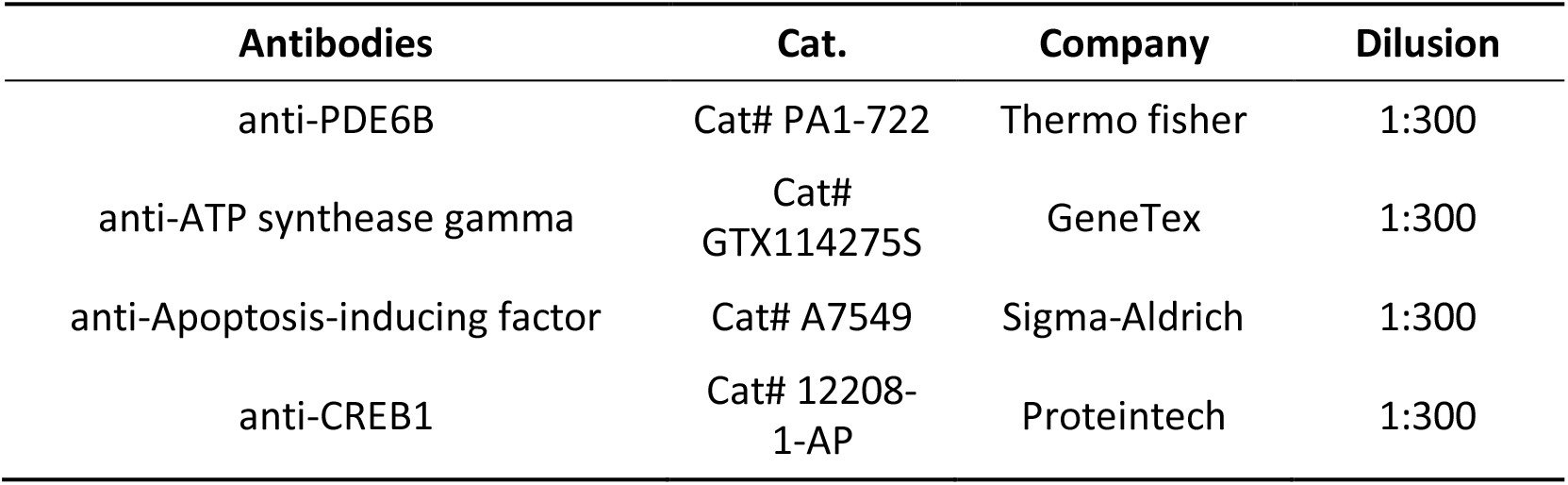
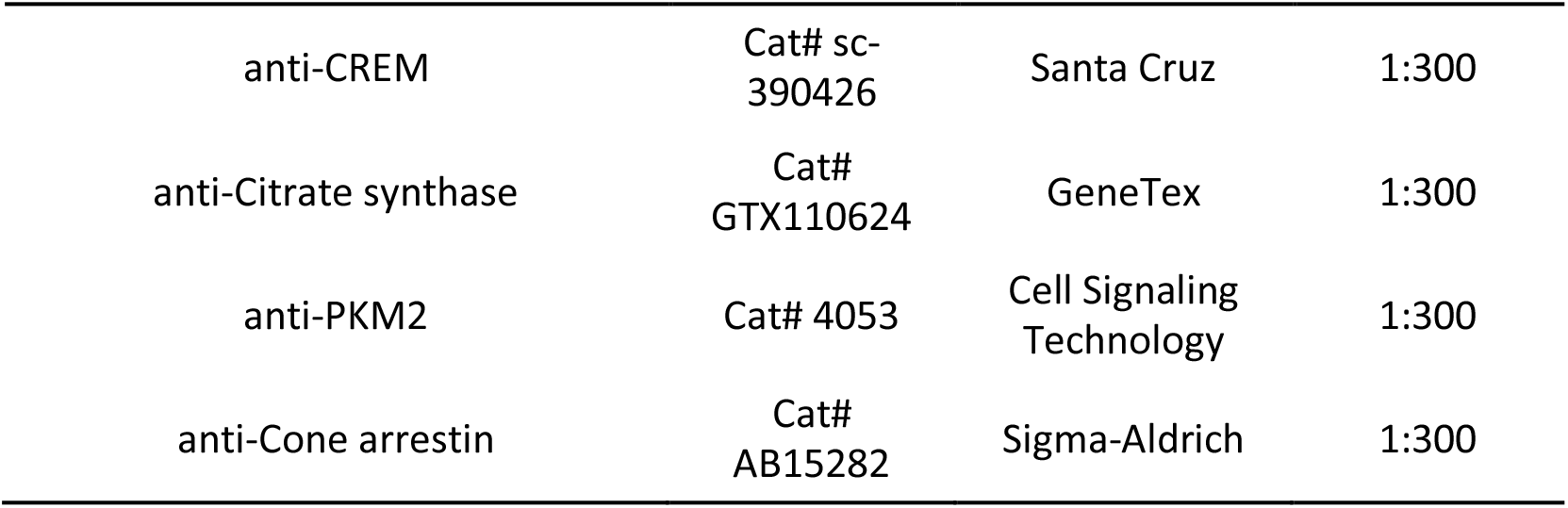
Primary antibodies used in the study.

### TUNEL assay

Terminal deoxynucleotidyl transferase dUTP nick end labelling (TUNEL) assay was performed using an *in situ* cell death detection kit (Fluorescein or TMR; Roche Diagnostics GmbH, Mannheim, Germany). Fixed slides were dried at 37°C for 30 min and rehydrated in phosphate-buffered saline (PBS) solution at RT, for 15 min. Afterwards, the slides were treated with proteinase K (Sigma-Aldrich Chemie GmbH, Taufkirchen, Germany) in TRIS buffer (10 mM TRIS-HCL, pH 7.4) at 37°C for 5 min. The slides were then washed with TRIS buffer three times for 5 min each. Subsequently, the slides were placed in ethanol-acetic acid mixture (70:30) at −20°C for 5 min followed by three washes in TRIS buffer and incubation in blocking solution (10% normal goat serum, 1% bovine serum albumin, 1% fish gelatin in 0.1% PBS-Triton X100) for 1 h at RT. Lastly, the slides were placed in the terminal dUTP-nick-end labelling (TUNEL) solution (labelling with either fluorescein or tetra-methyl-rhodamine) in 37°C for 1 h and cover slipped using Vectashield with DAPI (Vector, Burlingame, CA, USA).

### Tissue dissociation, single-cell preparation

WT and *rd1* mice at P11 (n = 3; Retina n = 6), P13 (n = 3; Retina n = 6), and P17 (n = 3; Retina n=6) were sacrificed regardless of gender. The eyeballs were quickly placed into DPBS (Dulbecco’s phosphate-buffered saline without Ca^2+^ and magnesium CAT:21-040-CVC, CORNING) pre-cooled at 4°C, incubated in 0.12% proteinase K (Millipore, 539480) at 37°C for 1 min and basal medium (Gibco, Paisley, UK) with 50% Foetal Bovine Serum (Gemini, 900-108) for 2 min, and then transferred to fresh DPBS for a final wash. Afterwards the cornea, sclera, iris, lens, and vitreous were removed on ice under the microscope, the retinal tissues containing retina-RPE-choroid were completely immersed in MACS tissue storage solution (Miltenyi, 130-100-008, Bergisch Gladbach, Germany) which was pre-cooled at 4°C and detected immediately to ensure that the activity and numbers of retinal cells were sufficient for further experimental analysis.

### Single-cell RNA-seq (scRNA-seq) and bioinformatics analysis

Retinal cellular suspensions (43,979 cells) were loaded on a 10x Genomics Chromium Single Cell instrument (10x Genomics, Shanghi, China) to generate single-cell Gel Beads in Emulsion (GEMs). Barcoded sequencing libraries were conducted following the instruction manual of the Chromium Single Cell 3’ Reagent Kits v3 (10x Genomics). Following the library preparation (Chromium Single Cell 3’ Reagent Kit v3 (10x Genomics (Shanghai) Co., Ltd, Shanghai, China), the sequencing was performed with paired-end sequencing of 150nt each end on one lane of Illumina NovaSeq 6000 (Illumina, San Diego, CA, USA) per sample. Retinal scRNA-seq analyses were performed using the Seurat package in R 14 (R Core Team (2022), R: A language and environment for statistical computing. R Foundation for Statistical Computing, Vienna, Austria. URL https://www.R-project.org/). Briefly, cells with a significant number of outlier genes (potential polysomes) and a high percentage of mitochondrial genes (potential dead cells) were excluded from using the “Filter Cells” function. The Log Normalize method was used to normalize gene expression. Principal component analysis (PCA) was then performed to reduce the dimensionality of the dataset using t-SNE/UMAP dimensionality reduction. Seurat was used to cluster cells based on the PCA scores. For every single cluster, differentially expressed genes (DEGs) were identified using the “Find All Markers” function in the Seurat package, and the screening threshold was set to |avg_logFC| > 0.58 and *p* < 0.05. The identified genes were functionally and taxonomically annotated with major databases, including the Gene Ontology (GO) enrichment analysis, the Kyoto Encyclopedia of Genes and Genomes (KEGG) and Metascape (https://metascape.org/), to understand the functional properties and classifications of different genes, and the results were shown by the R software (RStudio Team (2022). RStudio: Integrated Development for R. RStudio, PBC, Boston, MA URL http://www.rstu-dio.com/.).

## 5. Conclusions

The single-cell analysis described in this report provides transcriptional signatures for the diverse populations of neuronal and glial cells in the *rd1* mouse model for retinal degeneration. We focused on alterations in metabolic pathways in rod and cone photoreceptors in the critical time period from P11 to P17 during which most of the rod degeneration occurs. The identification of a metabolic switch likely related to altered activity of the MAPK-pathway highlights new possibilities for treatments designed to reprogram cellular metabolism so as to promote cell survival and recovery. Remarkably, the metabolism switch observed in *rd1* mutant rods may extend to genetically intact cones, suggesting that very similar treatments could rescue both rods and cones. On a broader level, our study provides further in-depth insights into the pathological mechanisms of rod-cone dystrophy.

## Supplementary Materials

The following supporting information can be downloaded at: www.mdpi.com/xxx/s1, Figure S1: title; Table S1: title; Video S1: title.

## Author Contributions

Conceptualization, K.J. and F.PD.; methodology, Y.C., W.L., S.Y., J.Y. and J.D.; writing—original draft preparation, Y.C. and J.Y.; writing—review and editing, Y.C., J.Y., K.J. and F.PD.; supervision, K.J. and F.PD.; funding acquisition, K.J. and F.PD..

## Institutional Review Board Statement

Protocols for animal experimentation were compliant with the German law on animal protection were reviewed and approved by the ‘Einrichtung fur Tierschutz, Tierärztlichen Dienst und Labortierkunde’ of the University of Tübingen, and registered under the number AK 02/19 M.

## Informed Consent Statement

Not applicable.

## Data Availability Statement

Our original single cell sequencing data is being reviewed by GEO.

## Acknowledgments

The author would like to thank Jiayin Biotechnology (Shanghai, China) for technical support.

