## Supplemental figures for "Single-cell transcriptomic profiling in inherited retinal degeneration reveals distinct metabolic pathways in rod and cone photoreceptors"

**Supplementary data and figures**

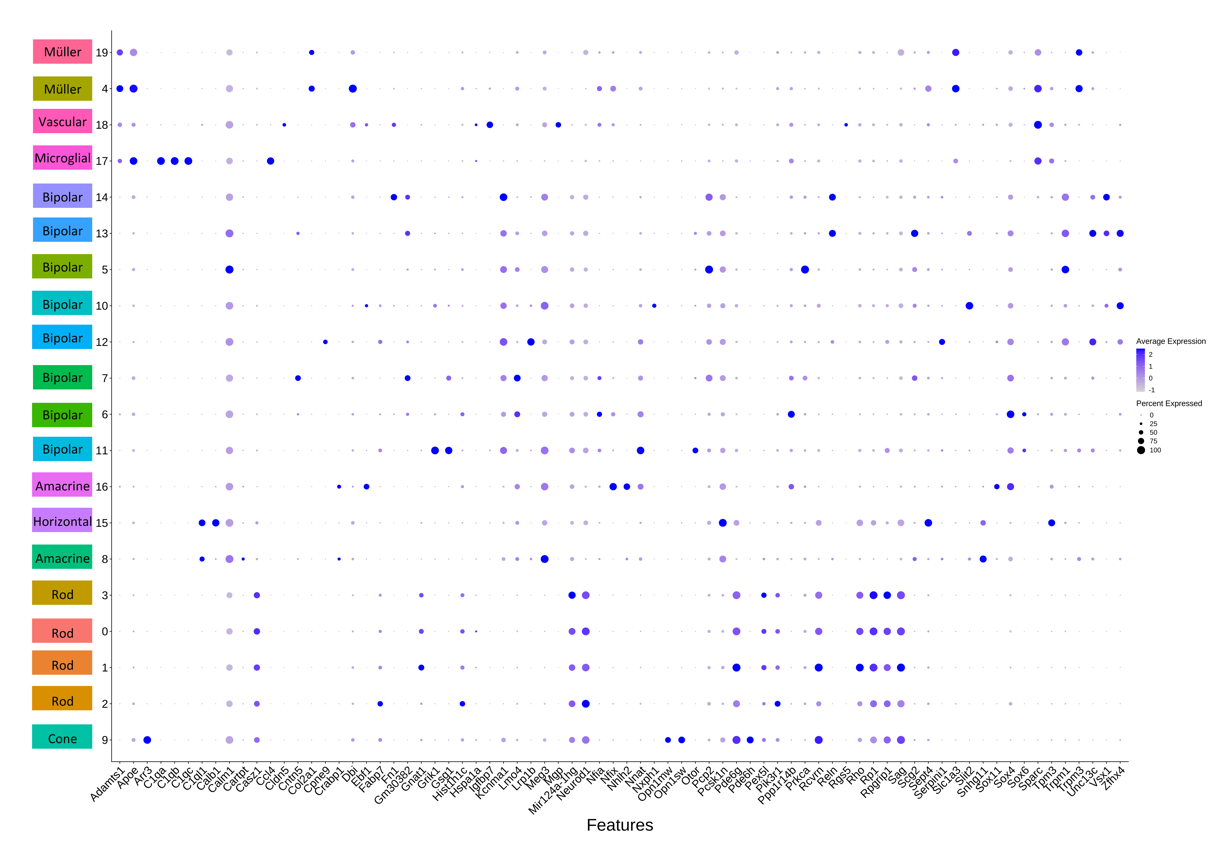

**Figure S1**. **Dot plot of the top four markers for each gene cluster.** The Seurat FindMaker function was used to analyze the differences between each cell population. Marker genes were selected based on an average log_2_ fold change (FC) > 1.5 and a *p*-value < 0.05. Genes are plotted on the x-axis, the y-axis indicates the cluster ID. A cell type related to a cluster is also shown on the left. The color ramp indicates the average expression of a certain gene in cell population. Dot size indicates the percentage of cells expressing the gene in that population.

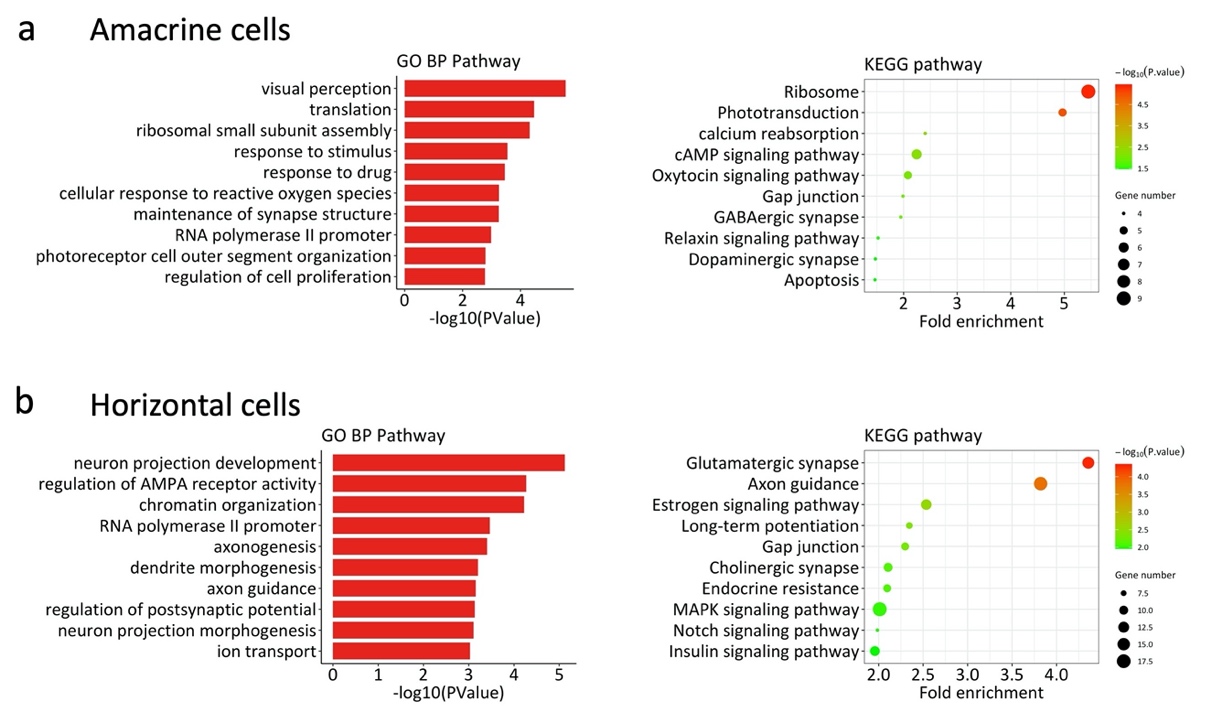

**Figure S2. Pathway analysis on amacrine and horizontal cells at post-natal day (P)13.** Data showing differentially enriched genes (DEGs) from *rd1* compared to wild-type (WT). Left panel: Gene ontology (GO) biological category (BP) analysis of the DEGs showing the top 10 most enriched GO BP terms. X-axis indicates statistical significance of differential expression as -log10 (*p*-value). Y-axis shows GO BP categories. Right panel: scatter plot of DEGs KEGG enrichment. X-axis indicates fold enrichment. Y-axis specifies KEGG pathways. Dot size indicates number of DEGs per pathway. Color coding indicates *p*-value range.

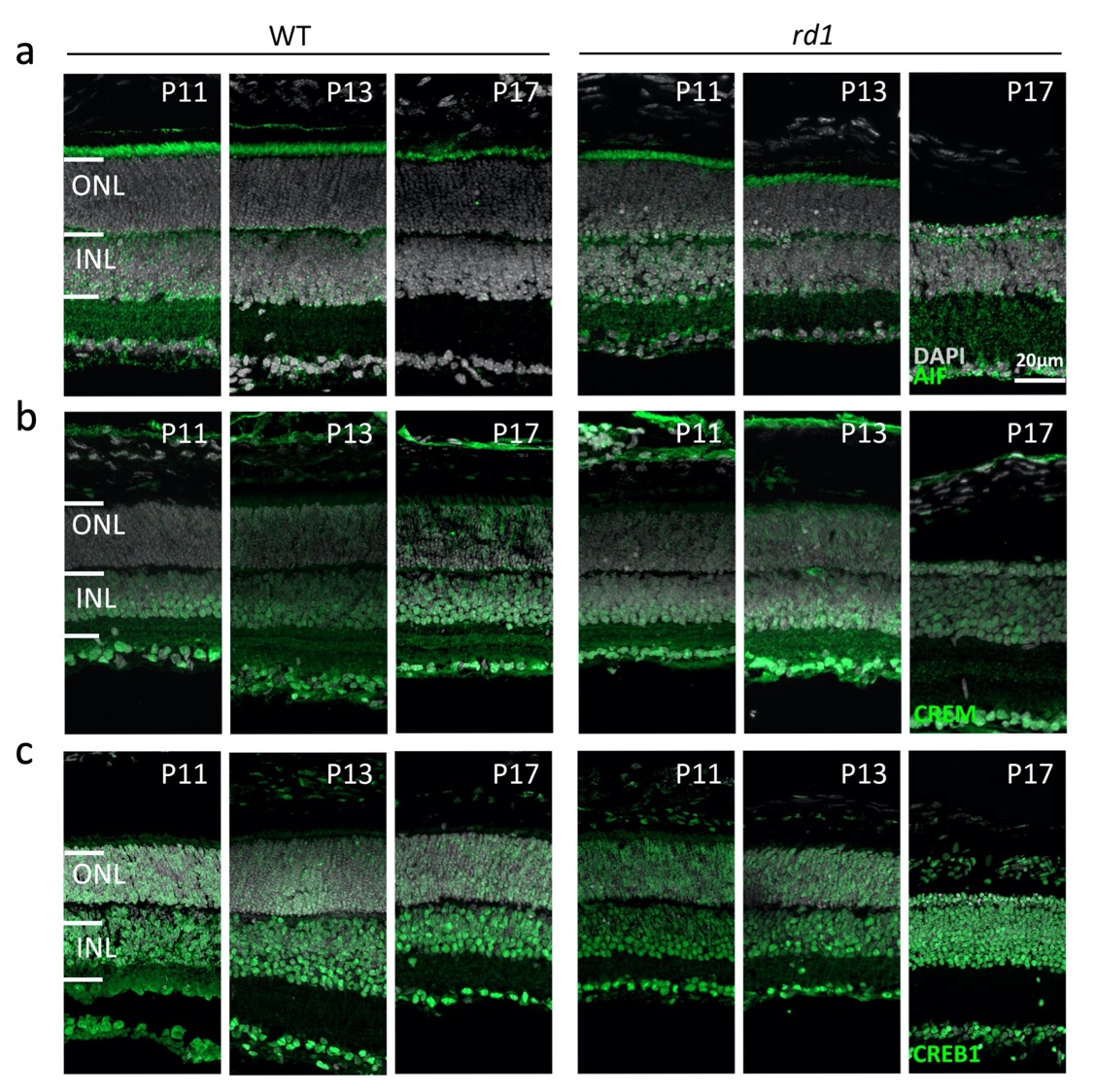

**Figure S3. Retinal expression of AIF, CREM, and CREB1 during the 2^nd^ post-natal week.** (**a**) AIF labelling of photoreceptor inner segments was mostly lost in P17 *rd1* retina. (**b**) IF for CREM labelled the entire retina, without obvious differences between WT and *rd1*. (**c**) Similarly, CREB1 labelling was seen throughout retina without apparent differences between WT and *rd1*. DAPI (grey) was employed as nuclear counterstain. ONL, outer nuclear layer; INL, inner nuclear layer; WT, wild-type; *rd1*, retinal degeneration 1.

**Supplementary dataset 1:** this dataset is an analysis of variance for comparing statistical differences in gene expression between cell populations of interest, which includes rods, cones, Müller glia cells, amacrine cells and horizontal cells. We use the Seurat and FindMarkers function of the R package for variance analysis to perform comparison between wild-type and *rd1* retinas. The screening threshold for differential genes was |avg_logFC| > 0.58 and p_val < 0.05. In dataset, feature: gene name. p_val: p-value of significance of differential expression. avg_logFC: log 2 fold change of the average expression value of the two cell populations. pct.1: the percentage of cells expressing the gene in the cell population of the experimental group. pct.2: percentage of cells expressing the gene in the cell population of the control group. p_val_adj: adjusted p-value. geneDescription: description of the gene.
